# Improving Targeting Specificity of Transcranial Focused Ultrasound in Humans Using a Random Array Transducer: A k-Wave Simulation Study

**DOI:** 10.1101/2025.04.25.650630

**Authors:** Zherui Li, Kai Yu, Joshua Kosnoff, Bin He

**Affiliations:** Department of Biomedical Engineering, Carnegie Mellon University, Pittsburgh, PA 15213, USA; Neuroscience Institute, Carnegie Mellon University, Pittsburgh, PA 15213, USA

**Keywords:** Transcranial Focused Ultrasound, Aberration Correction, Phase Reversal, V5, k-Wave, Computer Simulation

## Abstract

Transcranial focused ultrasound (tFUS) has emerged as a promising non-invasive modality for precision neuromodulation. However, the heterogeneous acoustic properties of the skull often induce phase aberrations that shift the ultrasound focus and compromise energy delivery. In this study, we developed and validated a phase-reversal based aberration correction method to enhance the targeting specificity of tFUS using a 128-element random array ultrasound transducer. Individual head models were constructed from T1-weighted magnetic resonance (MR) images and corresponding pseudo-computed tomography (pCT) data to accurately represent subject-specific skull geometries and the targeted left V5 (V5L) region. Acoustic simulations were conducted with the k-Wave toolbox by first acquiring free-field pressure waveforms and then recording the aberrated waveforms in the presence of the skull. The phase differences between these conditions were used to compute corrective delays for each transducer element. Quantitative evaluation using metrics such as focal overlap with the target region, axial focal positioning, and the delivered ultrasound energy demonstrated significant improvements: the overlap volume increased by 98.70%, mean axial positioning errors were reduced by up to 14.36%, and energy delivery to the target improved by 17.58%. We further demonstrated that the proposed approach outperforms the conventional ray-tracing methods. The results show that phase-reversal based aberration correction markedly increases the spatial targeting accuracy of tFUS and enhances the efficiency of focused ultrasound energy deposition for the customized random array transducer, paving a way for effective and personalized non-invasive neuromodulation therapies.

## 1. Introduction

Non-invasive neuromodulation techniques offer promising alternatives to invasive procedures, potentially reducing risks associated with surgical interventions and enhancing patient safety. It began with the advent of electroconvulsive therapy (ECT) in the early 20th century, which demonstrated that externally delivered electrical pulses could induce therapeutic seizures and alter neural excitability [1, 2]. Building on the principle that energy delivered non-invasively can reshape brain activity, in 1955, Fry *et al.* first demonstrated that high intensity focused ultrasound (HIFU) could precisely and safely ablate discrete targets in animal brain without damaging surrounding healthy tissues—establishing focused ultrasound as a spatially precise alternative to both ECT and surgical lesioning [3]. Recently, adapting that same focusing principle at much lower intensities, transcranial focused ultrasound (tFUS) has emerged as a powerful modality for modulating neural activities and treating brain disorders without invasive surgery [4, 5]. Specifically, low-intensity tFUS can either excite or inhibit neuronal activity by utilizing different acoustic parameters, providing a versatile tool for neuromodulation [6–13]. It can also be used for pain management, drug delivery, blood-brain barrier (BBB) opening, and tissue ablation/destruction under different sonication regimes through thermal or mechanical effects [14–22].

Traditional neuromodulation techniques each have their strengths, but they often come with limitations in either invasiveness or spatial resolution—challenges that tFUS effectively addresses. Deep brain stimulation (DBS), for instance, is an FDA approved and efficacious treatment for diseases like essential tremor (ET), Parkinson’s disease (PD), and epilepsy [23–25]. However, DBS and other similar invasive methods usually require surgical implantation of electrodes, posing risks of infection, bleeding, and other complications [26]. Non-invasive methods like transcranial magnetic stimulation (TMS) and transcranial direct current stimulation (tDCS) can also widely modulate superficial brain regions [2, 27], but with limited (multi-centimeter-scale) spatial resolution and deep brain penetration ability [28–30]. tFUS, on the other hand, enjoys high spatial precision (millimeter-scale), and can be steered to modulate shallow cortical areas or to penetrate deeper brain regions by modulating the source signal or using various transducers [6, 31]. However, the effectiveness of tFUS is significantly impacted by the skull’s complex structure, which introduces challenges such as ultrasound energy attenuation and phase aberration. Specifically, the varying density and irregular shape of the skull, as well as the significant acoustic property differences between the skull and soft tissues (e.g., the brain), can contribute heavily to ultrasound energy attenuation and phase aberration of the ultrasound waves [32, 33]. Such skull-induced phase aberration effect can shift and distort the ultrasound focus [32], making it difficult to deliver ultrasound energy to the target brain region. This not only reduces the efficacy of neuromodulation or treatment but also raises clinical concerns in ultrasound ablation procedures, as unintended brain areas could be targeted. It is essential to correct such phase aberrations by applying correction algorithms to tFUS transducers to maximize its efficacy, especially in clinical applications.

Prior studies on tFUS phase aberration correction have primarily focused on two key aspects: transcranial acoustic field estimation and transmitted pulse adjustment (through time reversal or phase compensation) [34]. For acoustic field estimation, the hydrophone method, which consists of directly recording pressures at the desired target, is currently regarded as the *gold standard*. However, it is not suitable for routine human applications. This is due to the potential for substantial brain tissue damage caused by implanting receivers into the cranial cavity or brain [35]. Though certain through-transmit-based approaches can further address this issue non-invasively by directly measuring aberrations with two transducers placed on opposite sides of the head, these approaches are unsuitable for pre-planning and remain limited by transducer orientations [36, 37]. Unlike such experimental methods, analytical methods and numerical methods are more practical for pre-treatment planning and neuromodulation purposes.

Among current analytical methods for transcranial acoustic field estimation, ray-tracing algorithms—such as those implemented in the open-source software Kranion— are commonly used approaches [38]. These algorithms rely on segmenting the skull and applying geometric approximations (based on Snell’s law) to determine the necessary phase shifts for each element in the transducer array [39–41]. These approaches are computationally efficient but may oversimplify the wave theory, especially in more complex regions where diffraction and scattering matter (e.g., superficial brain regions).

By contrast, numerical methods such as the finite-difference time-domain (FDTD) method and the k-space pseudospectral method can offer more accurate acoustic field estimation, though they may demand substantial computational resources [42–46]. For example, the k-space method implemented in the k-Wave toolbox used in this work can better capture the full physics of acoustic propagation, including diffraction, reflection, scattering, and absorption, even using a relatively coarse computational grid [47]. In addition, a common theme across previous works on such methods is the need for individualized patient data, i.e., computed tomography (CT) or magnetic resonance (MR) images [48, 49]. These data are used to account for spatial variations in skull thickness and acoustic properties (e.g., the speed of sound and tissue density) during acoustic field simulations. Highlighting the significance of these individual-specific simulations, Jing *et al.* further showed that after estimating the acoustic field with k-Wave simulations using a 1D linear array transducer and CT image, the transmitted ultrasound pulses can be adjusted using a time-reversal method to accurately correct phase aberrations, thus achieving tighter focal spots in an *ex-vivo* skull [43]. Although 1D arrays offer simple beamforming capabilities, they are limited in their ability to control focal steering in multiple dimensions and are less effective at compensating for skull-induced aberrations [50, 51]. Multi-element phased-array transducers, in contrast, provide greater flexibility in electronic beam steering and focusing, making them better suited for pre-planned or real-time phase correction in transcranial applications [52, 53].

To close this gap, we develop a phase-reversal aberration-correction approach designed to improve the focusing accuracy and ultrasound energy delivery performance of a 128-element random phased-array transducer (H-275, Sonic Concepts, Inc., Bothell, WA) when targeting the human left V5 (V5L, middle temporal visual area) in k-Wave simulations. Unlike previous studies focused on 1D or large-size phased arrays, our approach leverages the spatial flexibility of a smaller H-275 phased array to implement phase correction dynamically. To quantify how this correction improves both focus and energy delivery, we introduce four targeted evaluation metrics beyond the commonly used ones such as focal pressure and focal region size [34, 54, 55]: i) the overlap volume between the ultrasound focus and the target region (*V*_*overlap*_/mm3); ii) total ultrasound energy delivered to the target (*E*_*target*_/mJ); iii) axial mean position of the focal region (*X*_*focus*_/mm); iv) axial position of the focal peak (*X*_*peak*_/mm). Compared to cases without correction, the proposed correction approach can significantly increase *V*_*overlap*_ and *E*_*target*_, which suggests that the proposed approach can improve both ultrasound focusing and energy delivery performance. Meanwhile, the results after phase correction also show significant decreases of *X*_*focus*_ and *X*_*peak*_, suggesting the positioning error gets improved. Furthermore, this work provides a basis for subsequent tFUS-based neuromodulation and therapeutic studies on human V5 by enabling improved targeting specificity, particularly when employing phased-array transducers.

## 2. Methods

The goal of this study is to develop and validate a phase-reversal based aberration correction approach for tFUS. This approach aims to help precisely deliver ultrasound energy to specific brain targets despite skull-induced distortions. To achieve this, we first created individualized human head models, incorporating subject-specific skull and brain geometries. Then, acoustic simulations using the k-Wave toolbox were performed. Further, to address skull-induced ultrasound aberrations, a phase-reversal correction method was implemented. Finally, four quantitative metrics were introduced and comparative validations were conducted to assess focusing accuracy and energy delivery, allowing thorough evaluation of simulation results and the proposed correction approach.

### 2.1. Subject Recruitment

T1-weighted (T1w) magnetic resonance (MR) images were collected from 21 healthy human subjects (9 male/12 female; mean age: 22.65±3.98) as part of a previous tFUS experiment [56]. Subject recruitment and image collection complied with all relevant ethical regulations regarding human research and were reviewed and approved by the Advarra Institutional Review Board (protocol number: STUDY2017_00000426). Additionally, a standard head model template (MNI-152) was also employed as a baseline reference during pre-experimental analysis [57–59]. Thus, a total of 22 human head models were constructed and utilized in this work.

### 2.2. Construction of Individualized Head Models for Acoustic Simulations

Each individualized human head model contained two parts in space, i.e., the target brain region of interest (ROI; V5L) and the skull bone (Figure 1a). In primates, V5 contains direction-selective neurons and plays a vital role in motion perception as well as integration of local motion signals into global percepts [60]. Given such functional characteristics, we selected left V5 (V5L) as the ROI because precisely targeted neuromodulation of this region could significantly impact visual perception research and therapeutic interventions requiring modulation of visual processing.

**Figure 1.**
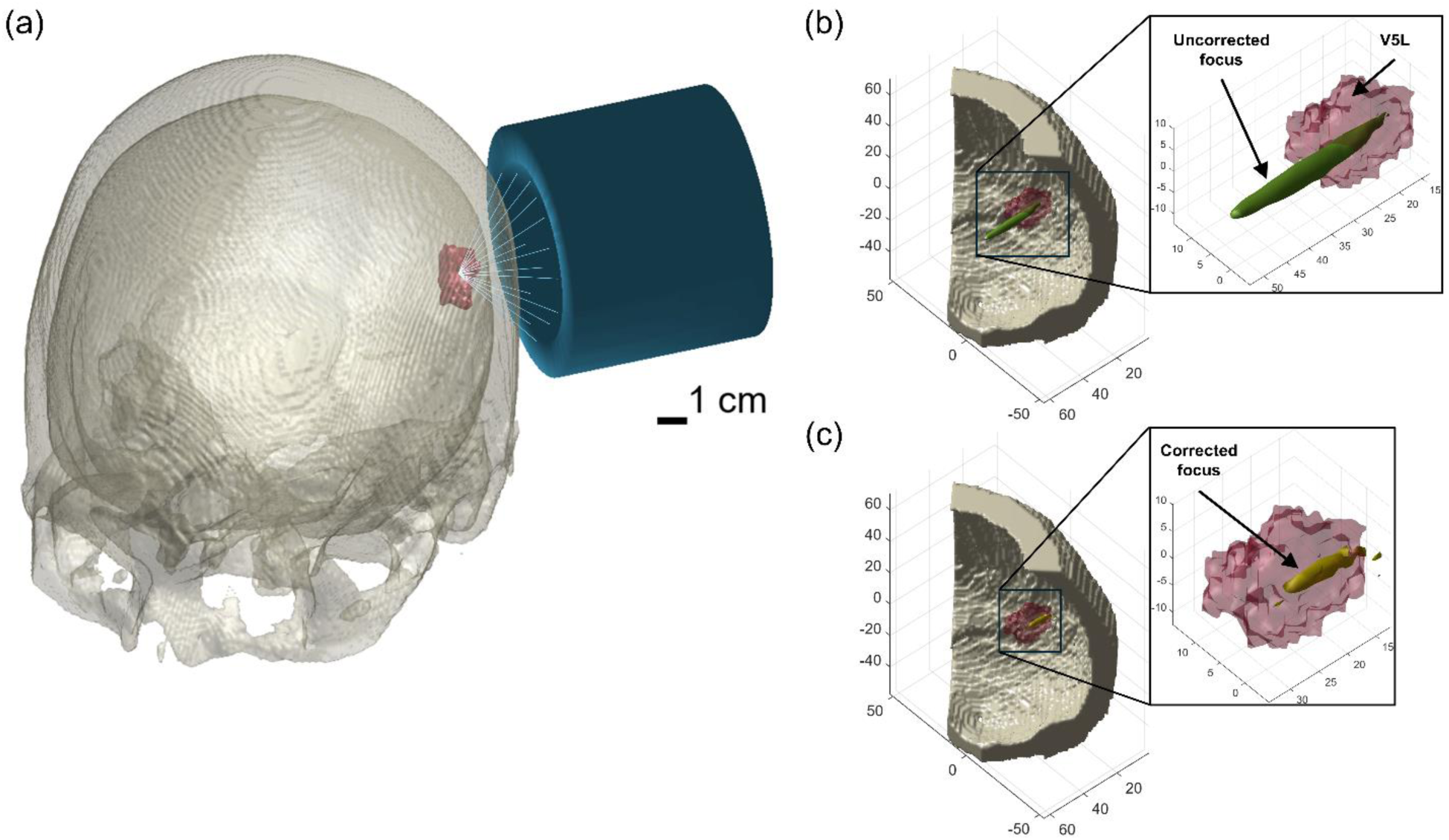
Schematic diagram of the approach. (a) Individualized 3D human head model is constructed from MR and pCT images, which includes the skull bone (bone-colored, semi-transparent) and the V5L region (red). The H-275 ultrasound transducer (blue) is aligned to target V5L. (b) and (c) compare the ultrasound focal region before and after applying the phase aberration correction, demonstrating that the proposed method effectively improves tFUS focusing on V5L (skull: bone-colored; V5L: red, semi-transparent; ultrasound focus: green/yellow; unit of length along each axis: mm).

To segment V5L for each subject, we developed a five-step protocol using the FMRIB Software Library (FSL) [61]: i) *brain extraction* from T1w MR images with BET (Brain Extraction Tool) [62]; ii) *linear registration* to the MNI-152 standard space with FLIRT (FMRIB’s Linear Image Registration Tool) [63]; iii) *non-linear registration* with FNIRT (FMRIB’s Nonlinear Image Registration Tool) to better accommodate differences in the shape of brain structures between the subject and the standard space [64]; iv) *extract and save* V5L region in the standard space from the Juelich structural atlas as a mask [65]; v) *invert the transformations* from the standard space to subject space and *apply the inverse warp* to the mask, which generates a native V5L mask in subject space. It should be noted that such V5L masks generated through this protocol were binary masks for post-simulation focus evaluations only and they were not involved in k-Wave simulations. The skull model of each subject was developed by thresholding the pseudo-CT (pCT) images generated from the subject’s T1w MR images. The threshold value for each subject’s pCT ranged from 70 to 140 to clearly define contours of the skull and to eliminate the artifacts within the cranial cavity. In addition, we adopted an open-source toolbox developed by Yaakub *et al.* to generate such pCT images [66]. This toolbox was utilized since it was specifically designed for acoustic simulations of transcranial ultrasound stimulation (TUS) and was able to let us avoid exposing subjects to unnecessary ionizing radiation due to additional CT scans. All generated pCT images were visually inspected to confirm their quality before thresholding. It should be noted that all skull models are also binary masks in space to reduce the error in Hounsfield units (HU) caused by MR to pCT conversion and thus the accuracy of density and speed of sound (SOS) calculations in later simulations, as well as to reduce the consumption of computational resources.

### 2.3. Acoustic Simulation Setup

In k-Wave simulations, the computational space was 70 mm in the axial direction (x-axis) and 150 mm in the lateral direction (y- and z-axis), which gave an input grid of 86×140×140 voxels with an isotropic spacing (*dx*) of 1.07 mm. In addition, a perfectly matched layer (PML) was applied, by setting its size to auto, to allow effective absorption of the ultrasound energy at grid boundaries and to avoid the reflection of ultrasound waves. Consequently, this offered an expended computational grid size of 128×162×162 voxels, which can well include all necessary components for each simulation: the ultrasound transducer (Figure 1a) and a piece of skull over the left occipital lobe (to increase the spatial resolution at the focal area) that covers the V5L region (Figure 1b and Figure 1c).

Specifically, the ultrasound transducer, implemented as 128 disc-shaped elements in simulations, was fixed at the same location in space. The spatial location and orientation of each element were defined according to the physical design of the H-275 transducer. Then, by visually targeting the spatial center of the V5L for each subject, the masks of skull and V5L were shifted and rotated accordingly.

Furthermore, considering that the skull significantly contributes to ultrasound aberration, soft tissues were excluded from the simulations. Acoustic parameters for the skull and surrounding water medium were then defined based on established literature [67, 68]: the skull properties were set with a speed of sound (*c*_*skull*_) of 2850 m/s, density (*ρ*_*skull*_) of 1732 kg/m^3^, and attenuation coefficient (α_*skull*_) of 85 Np/m/MHz. The surrounding water medium properties were set with a speed of sound (*c*_*water*_) of 1482 m/s, density (*ρ*_*water*_) of 1000 kg/m^3^, and attenuation coefficient (α_*water*_) of 3.48×10^-4^ Np/m/MHz. Then, to maintain numerical stability and computational efficiency, simulations employed a Courant-Friedrichs-Lewy (CFL) number of 0.02. The total computation duration (*t*_*end*_) of 50 μs allowed sufficient time for wave propagation and convergence. Additionally, the source frequency (*f*_0_) was set to 700 kHz to match the central frequency of the physical H-275 transducer, with an amplitude (*S*_*amp*_) of 1 MPa on an element basis. Notably, such disc-shaped elements with a diameter of 4 mm each were defined in a kWaveArray class, with a focal depth set to 35 mm (matching with that of the physical transducer).

### 2.4. Aberration Correction by Phase Reversal

With the head model and acoustic setup in place, we developed a wave theory-informed k-Wave based phase-reversal method to tackle skull-induced aberrations (Figure 2). Initially, each transducer element was individually activated and fired in a free field (i.e., only water as medium in the computational grid) with a series of continuous waves (CW) as source. The CW signal was defined based on *t*_*end*_, *f*_0_, *S*_*amp*_, and a phase of 0. Then, record the pressure at the intended target location (i.e., the physical focus of the random array transducer) to obtain 128 free-field waveforms. Such waveforms contain the reference phase information for aberration correction.

**Figure 2.**
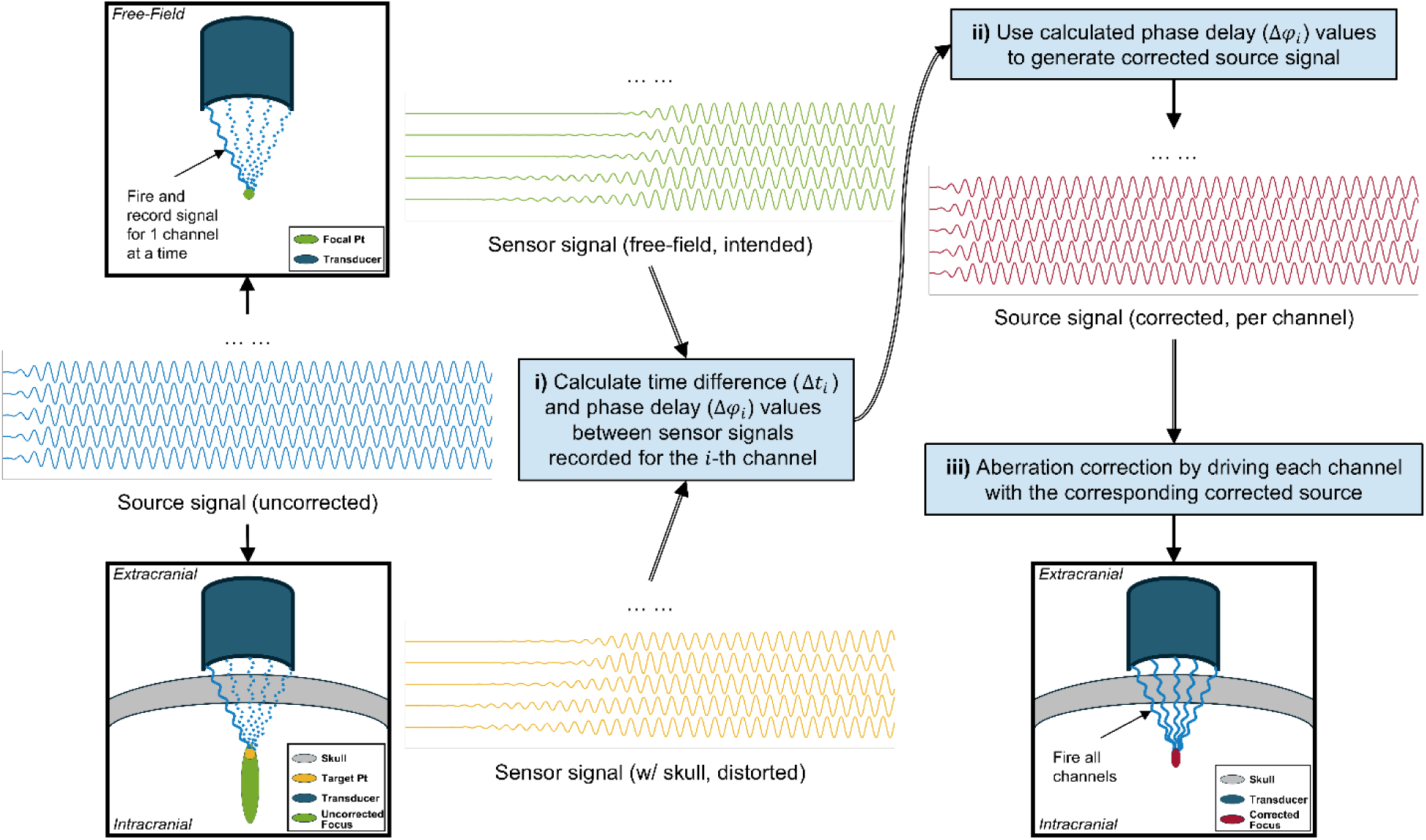
The workflow of the proposed phase aberration correction method for a random array transducer. i) A continuous wave (CW) signal without phase modulation (waveforms in blue) is sequentially emitted from 128 channels in free-field and skull-present conditions. The signals received at both the intended focal point (waveforms in green) and the desired target point (waveforms in yellow) are recorded. For the i-th (i ∈ [1,128]) channel, peak detection is then performed to compute the time difference (Δ*t*_*i*_) based on the last detected peak pair, allowing for phase difference (Δφ_*i*_) estimation; ii) The estimated phase differences for each channel are then used to generate corrected CW source (waveforms in red); iii) The corrected source will be further used to drive their respective channels, which can ensure phase aberration correction at the desired target point and thus constructing a corrected focus.

Subsequently, the skull was introduced into the simulation grid. Then, the procedure is repeated (i.e., firing each element with the same CW signal), recording new waveforms distorted by the skull-induced aberration at the intended location. Finally, the time difference (Δ*t*_*i*_) between the *i* -th pair of waveforms (*i* ∈ [1,128]) recorded with and without the skull can be calculated through Equation 1 based on the last pair of peaks, where *maxl*_*free*−*field,i*_(*end*) and *maxl*_*skull*,*i*_(*end*) correspond to the last time steps where we can find a peak from the recorded waveforms, respectively. Then, the phase delay values (Δφ_*i*_) for the *i*-th transducer element can be calculated by Equation 2, where δ*t* is the temporal spacing of the simulation calculated with Equation 3 with a points-per-wavelength (PPW) value set to 2.

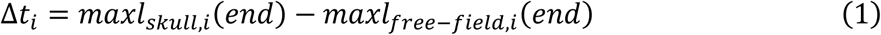

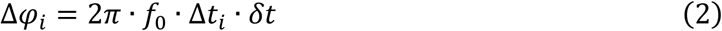

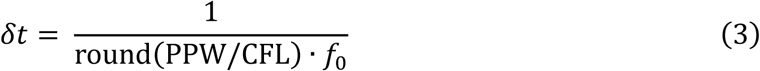

By incorporating these calculated Δφ_*i*_ into the CW source signals for each transducer element accordingly, the skull-induced aberrations can be effectively corrected (Figure 1b and Figure 1c), enhancing the precision and efficacy of the tFUS focusing procedure.

### 2.5. Quantitative Evaluation Metrics

To demonstrate how effectively the proposed method corrects skull-induced aberrations, two main groups of quantitative metrics were introduced: focusing metrics and energy delivery metrics. Focusing metrics were only applied to the axial direction since the skull-induced aberrations dominantly affect the location and shape of the focus axially rather than laterally according to our simulations (Table I).

**Table I.** Quantitative Comparison of tFUS Focus Position and Spatial Resolution with and without Aberration Correction (*N* = 22)

| | $X_{focus}/\text{mm}$ | $Y_{focus}/\text{mm}$ | $Z_{focus}/\text{mm}$ | $d_x/\text{mm}$ | $d_y/\text{mm}$ | $d_z/\text{mm}$ | $X_{peak}/\text{mm}$ |
| --- | --- | --- | --- | --- | --- | --- | --- |
| No Correction | $38.37 \pm 2.66$ | $0.00 \pm 0.00$ | $0.00 \pm 0.00$ | $31.38 \pm 3.22$ | $3.57 \pm 0.71$ | $3.62 \pm 0.54$ | $31.93 \pm 6.30$ |
| Corrected | $32.86 \pm 7.01$ | $0.00 \pm 0.48$ | $-0.15 \pm 0.38$ | $32.19 \pm 12.68$ | $3.57 \pm 1.04$ | $4.08 \pm 1.42$ | $25.05 \pm 8.63$ |

#### 2.5.1. Focusing Metrics

To measure the focusing performance between the scenarios with and without aberration correction, both the axial location of the focus and the overlap volume between the focus and the target are worth considering. Therefore, we introduced the axial mean position of the focal region (*X*_*focus*_/mm), the axial position of the focal peak (*X*_*peak*_/mm), and the overlap volume between the ultrasound focus and the target region (V5L) (*V*_*overlap*_/mm^3^). To measure such metrics, the focus was first defined by -6 dB (full width at half maximum) relative to the peak intracranial pressure in space, which is a commonly used approach for focal region definition [21]. *X*_*focus*_ was defined by the difference of the maximum and minimum axial values of the focus while *X*_*peak*_ corresponds to the axial location of the peak intracranial pressure. *V*_*overlap*_ was calculated from the overlap voxels of the focus and the V5L mask.

#### 2.5.2. Energy Delivery Metrics

Due to the varying nature of the ultrasound pressure at focus and the underlying mechanism of tFUS neuromodulation (thermal or mechanical effects induced by ultrasound energy) [69], considering only the ultrasound pressure or intensity may well quantify energy distribution of the focus itself [70], but is not sufficient for evaluating the fundamental question, “How much ultrasound energy is effectively delivered to the intended target region for neuromodulatory effects?” So, the total ultrasound energy delivered to the target (*E*_*target*_ /mJ) was introduced to answer this critical question. Derivation of *E*_*target*_ from the recorded pressure waveforms and *V*_*overlap*_ is further discussed below.

The acoustic energy density (*w*) in a particular time *t* and position <u>*E*</u> can be defined as the sum of the acoustic kinetic energy density (*w*_*v*_) and the acoustic potential energy density (*w*_*p*_) [71]:

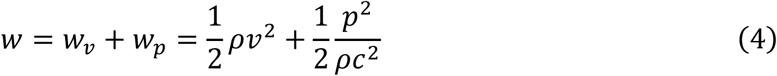

where *v* is the velocity of the acoustic wave, *ρ* is the density of the medium, and *c* is the speed of sound (SOS) in the medium. As intracranial soft tissues (e.g., brain tissues) are not included in the simulation, we take water as the medium here, i.e., *ρ* = *ρ*_*water*_ and *c* = *c*_*water*_.

Since the acoustic intensity (*I*) has such a relationship with *w*:

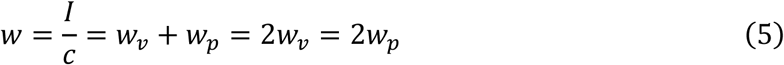

we can have:

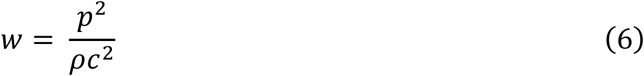

It should be noted that Equation 6 also holds when time varies, i.e., 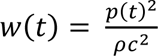 as *w*_*p*_ and *w*_*p*_ are integrated only over the volume [72]. In our simulations, CW (i.e., a series of sine waves) was used as the source signal as mentioned earlier. So, the instantaneous pressure *p*(*t*) varies sinusoidally with time:

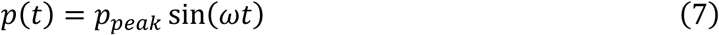

where *p*_*peak*_ is the peak pressure amplitude and ω is the angular frequency. Since that RMS (root mean square) pressure (*p*_*rms*_) represents time-averaged energy and for sine waves it can be given by:

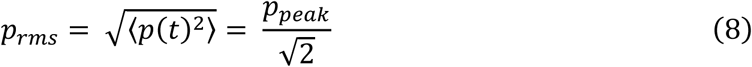

where 〈⋅〉 denotes the time average over one period of oscillation. Then, we can calculate the time-averaged acoustic energy density (*E*_*density*_) by [73, 74]:

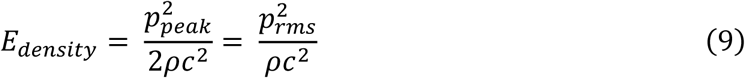

Since *E*_*density*_ only gives the energy per unit volume while we want to calculate the total energy delivered into the overlap region (i.e., *E*_*target*_), it can be simply defined by:

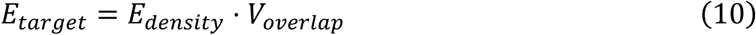

### 2.6. Additional Validation of Efficacy of the Proposed Aberration Correction Method Using the Ray-Based Kranion Software

To provide essential baseline validation and additional insights into the performance of the proposed aberration correction method, a comparative analysis using a ray-based software Kranion was done [38]. Specifically, the same human head models (i.e., skulls in pCT images with corresponding V5L in MR images) were imported to Kranion, in which the H-275 array transducer was also modeled with the same relative spatial location and orientation for each of its 128 elements. Due to time and computational resource constraints, we only randomly selected 11 subjects for such Kranion simulations (*N* = 11). In each Kranion simulation, the estimated phase delay values and element activation criterion (i.e., the estimated amplitude in 0 or 1) were saved into single file. Later, we applied them in additional k-Wave simulations separately or together for better comparison with the existing data obtained without correction or with our correction method. For the purposes of labeling such four cases later, we refer to the case without correction as *No Corr*, the case with our k-Wave based phase-reversal method as *kPR*, the case with element activation criterion generated by Kranion alone (phase delay estimated by our method) as *KE*, and the case with both element activation criterion and phase delay generated by Kranion as *KE+Ph*.

### 2.7. Software and Computational Resources

MATLAB (R2024b, The MathWorks Inc., Natick, Massachusetts) and k-Wave MATLAB Toolbox (version 1.4) were used for coding and acoustic simulations [47, 75]. In addition, all 3D k-Wave simulations were performed on a workstation equipped with an NVIDIA RTX 4090 GPU and a GPU server equipped with eight NVIDIA RTX A5000 GPUs.

## 3. Results

Through comprehensive simulations, we first demonstrate the proposed phase-reversal based aberration correction method’s effectiveness in significantly improving the positioning accuracy of the tFUS focus (mean axial positioning errors can be reduced by 14.36% and focal peak positioning errors can be reduced by 21.53%). Subsequently, we examine how these improvements translate into enhanced targeting specificity (the overlap volume increases by 98.70%) and energy delivery efficiency (energy delivery increases by 17.58%) to the designated brain region (i.e., V5L), emphasizing the proposed method’s practical value for tFUS applications. Furthermore, we statistically compare the correction performance using Kranion, which shows that ray-based methods may not well address the skull-induced aberration, whereas the proposed method outperforms it across all evaluation metrics.

### 3.1. Aberration Correction Significantly Improves tFUS Focus Position

The primary objective of aberration correction is to restore the spatial position of the ultrasound focus to its ideal configuration. In free-field simulations, the transducer array successfully generates a highly compact focal region (with an axial resolution of 5.36 mm and lateral resolution of 1.07 mm) precisely located at the expected axial position of 18.21 mm and lateral position of 0.00 mm (Figure 3a). This ideal scenario serves as the baseline for evaluating the effectiveness of the proposed aberration correction method.

**Figure 3.**
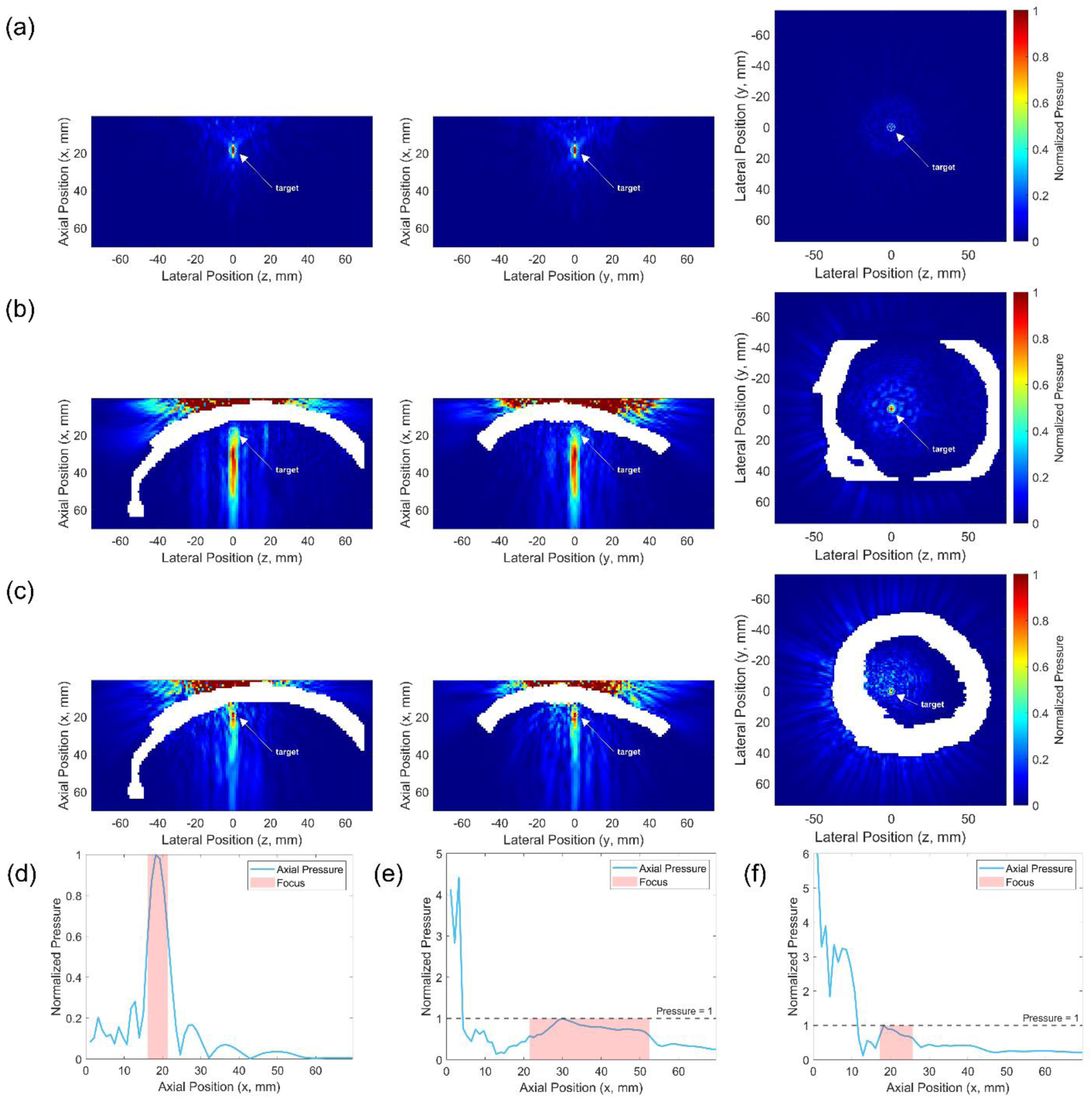
Simulation results. (a) Cross-sections of the simulated acoustic pressure field in a free field (left: coronal; middle: sagittal; right: axial). A highly compact focus is observed at the physical focal location of the transducer array (i.e., [*x*, *y*, *z*] = [18.21,0,0] mm). (b) Cross-sections of the simulated pressure field without phase aberration correction. While the focal region does not expand significantly laterally, it is notably elongated along the axial direction. Additionally, the focal peak shifts axially to [30,0,0] mm, misaligned from the intended target. (c) Cross-sections of the simulated pressure field after applying the proposed phase aberration correction method. The correction significantly improves the axial position and shape (i.e., axial resolution) of the ultrasound focus, realigning the focal peak with the physical focus, as expected. (d), (e), and (f) present the pressure profiles along the focal axis (*y* = 0) in the sagittal plane for the three conditions described above. Again, the proposed correction method effectively enhances the axial position and resolution of the ultrasound focus. However, significant pressure loss is observed due to skull-induced reflections and other aberration effects. *Note: When in the presence of a skull, the pressure normalization is performed relative to the maximum intracranial pressure value*.

In contrast to this idealized condition, the presence of the skull significantly reduces intracranial ultrasound focus resolution and causes an axial displacement of the focal region away from its expected position (Figure 3b). Since the mean lateral positions of the focal region (calculated at -6 dB) did not deviate from the expected location across all subjects significantly (Table I), we focused our analysis exclusively on axial effects caused by the skull, particularly the mean axial position of the focal region and the axial location of its peak.

After applying the proposed aberration correction method, both the axial resolution and axial positioning of the ultrasound focus notably improved for the same subject (Figure 3c). When considering the axial mean position of the focal region (*X*_*focus*_) and the axial position of the focal peak (*X*_*peak*_) for 22 subjects, significant improvements were observed (Figure 4a and Figure 4b). Compared to the uncorrected condition, the proposed method reduced *X*_*focus*_ by an average of 14.36% (from 38.37±2.66 mm to 32.86±7.01 mm, *p* = 3.09×10^-4^) and *X*_*peak*_ by 21.53% (from 31.93±6.30 mm to 25.05±8.63 mm, *p* = 2.63×10^-4^), bringing both values substantially closer to their respective ideal free-field values (19.29 mm for *X*_*focus*_ and 18.21 mm for *X*_*peak*_). Note that significance analyses here and below were performed using two-tailed paired t-tests.

**Figure 4.**
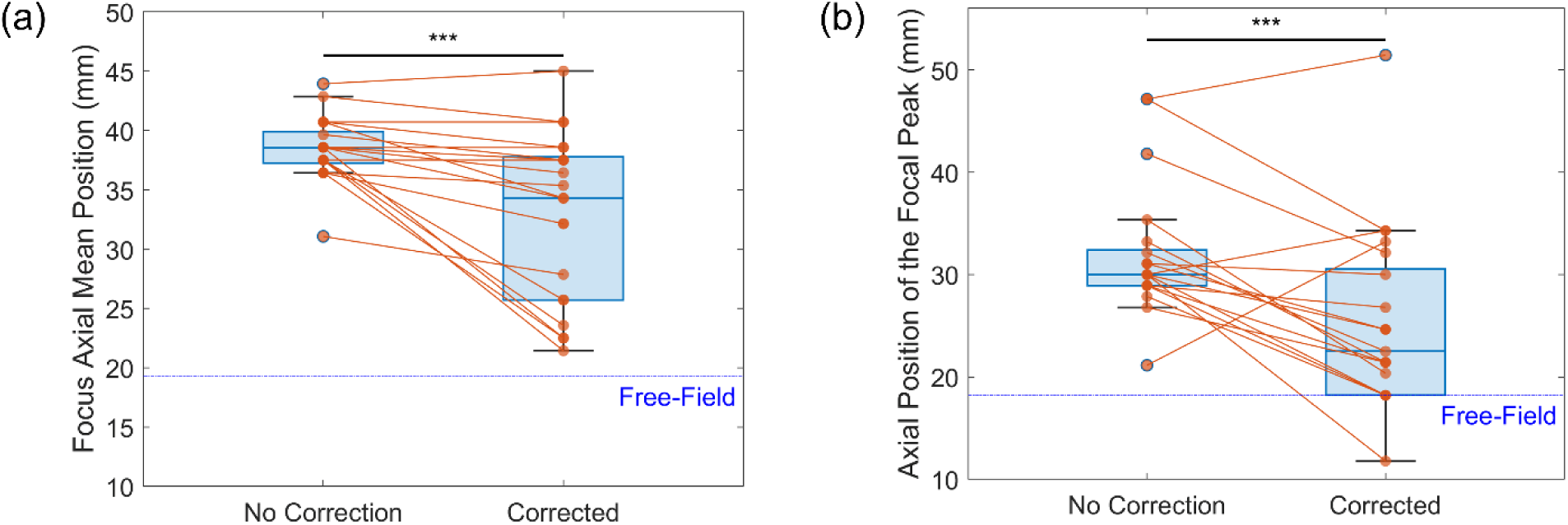
Quantitative measurements on the k-Wave simulation results in terms of focus positioning accuracy (*N* = 22. (a) Axial mean position of the focal region (*X*_*focus*_ /mm) decreased by 14.36%, from 38.37±2.66 mm in the uncorrected case to 32.86±7.01 mm after correction (*p* = 3.09×10^-4^). Additionally, the corrected position is closer to the ideal free-field value (x = 19.29 mm). (b) Axial position of the focal peak (*X*_*peak*_ /mm) decreased by 21.53%, from 31.93±6.30 mm in the uncorrected case to 25.05±8.63 mm after correction (*p* = 2.63×10^-4^). Notably, for several subjects, this metric reached the ideal free-field value (x = 18.21 mm) after applying the proposed correction method. *Note: Significance analysis was assessed using a two-tailed paired t-test; Key: ***p<0.001*.

A comparison of all metrics related to the spatial positioning and resolution of the ultrasound focus with and without aberration correction is presented in Table I, where *Y*_*focus*_ and *Z*_*focus*_ represent the lateral mean position of the focal region along the y- and z-axis, respectively. Meanwhile, *d*_*x*_, *d*_*y*_, and *d*_*z*_ represent the diameter of the focal region along the x-, y-, and z-axis, respectively.

### 3.2. Aberration Correction Enhances tFUS Targeting Specificity and Ultrasound Energy Delivery to V5L

Improving only the spatial position of the ultrasound focus itself is insufficient for tFUS-based neuromodulation or other applications. The ultimate goal is to effectively deliver ultrasound energy generated by the transducer to a specific target region or brain area to achieve desired modulatory or therapeutic effects via thermal or mechanical mechanisms. Therefore, aberration correction should also effectively increase the overlap volume (*V*_*overlap*_) between the focus and target region (V5L in this study) and enhance the total ultrasound energy delivered to this region (*E*_*target*_).

Through simulations (*N* = 22), the proposed aberration correction method significantly increased both *V*_*overlap*_ (Figure 5a) and *E*_*target*_ (Figure 5b). Specifically, the correction increased *V*_*overlap*_ by an average of 98.70% (from 51.58±18.80 mm^3^ to 102.49±38.12 mm^3^, *p* = 2.65×10^-7^) and *E*_*target*_ by 17.58% (from 6.07±3.08 mJ to 7.14±2.92 mJ, *p* = 1.30×10^-2^). Such improvements indicate that the proposed aberration correction method effectively enhances target coverage, thus reducing potential unintended effects on non-target brain regions. Furthermore, it demonstrates increased efficiency in energy delivery, which can be critical for the efficacy of tFUS applications.

**Figure 5.**
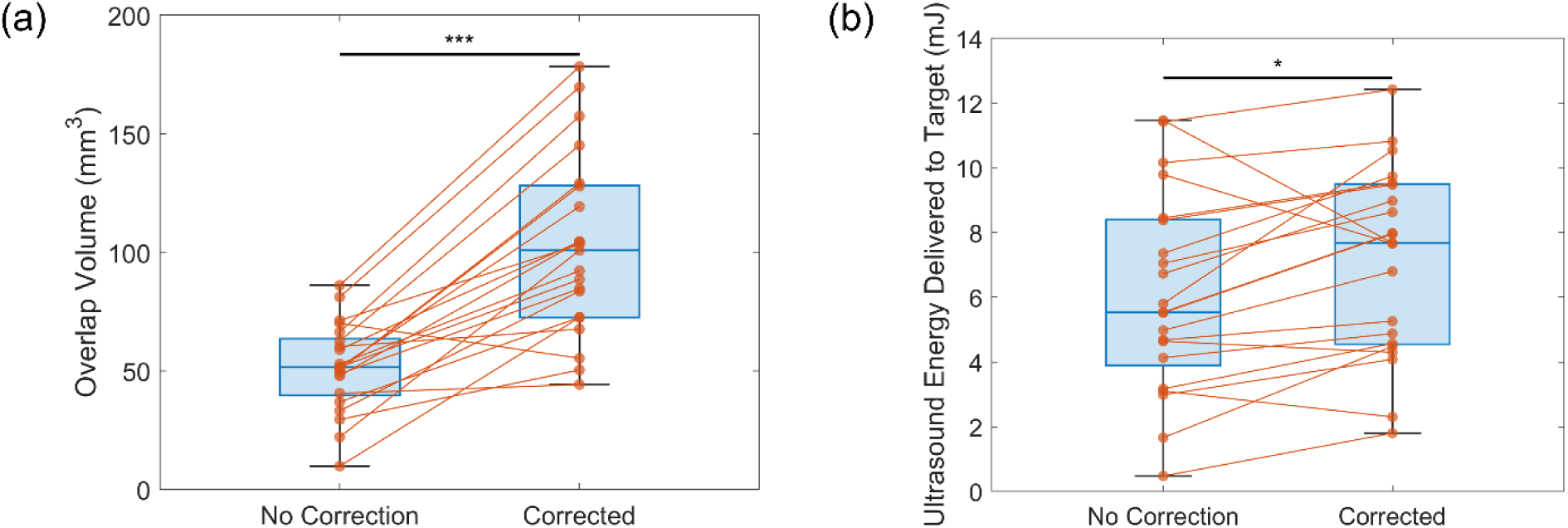
Quantitative measurements on the k-Wave simulation results in terms of targeting specificity and ultrasound energy delivery(*N* = 22. (a) Overlap volume between the ultrasound focus and the target region (V5L) (*V*_*overlap*_ /mm^3^) increased significantly from 51.58±18.80 mm^3^ in the uncorrected case to 102.49±38.12 mm^3^ after phase aberration correction, representing a 98.70% improvement (*p* = 2.65×10^-7^). (b) Total ultrasound energy delivered to the target (*E*_*target*_/mJ) increased from 6.07±3.08 mJ in the uncorrected case to 7.14±2.92 mJ after correction, with an improvement of 17.58% (*p* = 1.30×10^-2^). *Note: Significance analysis was assessed using a two-tailed paired t-test; Key: *p<0.05, ***p<0.001*.

Detailed comparisons of all V5L-related metrics are also provided in Table II. Note that the overlap rate was calculated relative to the volume of V5L (*V*_*V*5*L*_):

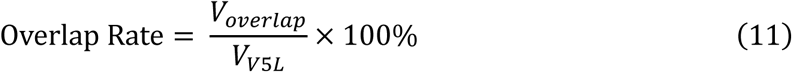

**Table II.** Quantitative Comparison of Target Region Overlap and Ultrasound Energy Delivery to V5L with and without Aberration Correction (*N* = 22)

| | $V_{overlap}/\text{mm}^3$ | Overlap Rate/% | $E_{target}/\text{mJ}$ |
| --- | --- | --- | --- |
| No Correction | 51.58±18.80 | 3.87±5.16 | 6.07±3.08 |
| Corrected | 102.49±38.12 | 8.25±13.22 | 7.14±2.92 |

### 3.3. The Proposed Phase-Reversal Aberration Correction Method Outperforms Kranion’s Ray-Based Method

For *V*_*overlap*_ (Figure 6a), while KE+Ph (i.e., both phase delay estimation and element activation criterion by Kranion) achieved the highest value of 133.84±45.12 mm^3^, this metric alone proved insufficient information for fully assessing the location or shape of the focal region. To address this, we first analyzed its *X*_*focus*_ (Figure 6c), *X*_*peak*_ (Figure 6d), and *d*_*x*_ (Table III). It can be observed that KE+Ph has a significantly lower *X*_*focus*_(36.33±2.60 mm) compared to that of No Corr (i.e., no aberration correction; 37.99±1.30 mm, *p* = 4.27×10^-2^). Then, for *X*_*peak*_, KE+Ph (29.81±8.86 mm) has a larger value than No Corr (29.10±3.13 mm, *p* = 0.81) but without group level significance. However, KE+Ph (40.62±4.51 mm) has a significantly larger *d*_*x*_ than No Corr (31.36±0.84 mm, *p* = 6.94×10^-5^). Thus, KE+Ph produced similar results compared to No Corr in terms of axial positioning but with worse focal shape (e.g., elongated along the axial axis). This effect can be further verified by their focal volume (*V*_*focus*_/mm^3^) through additional calculations, i.e., KE+Ph (628.28±320.22 mm^3^) has a significantly larger *V*_*focus*_ compared to No Corr (316.44±16.69 mm^3^, *p* = 7.79×10^-3^).

**Figure 6.**
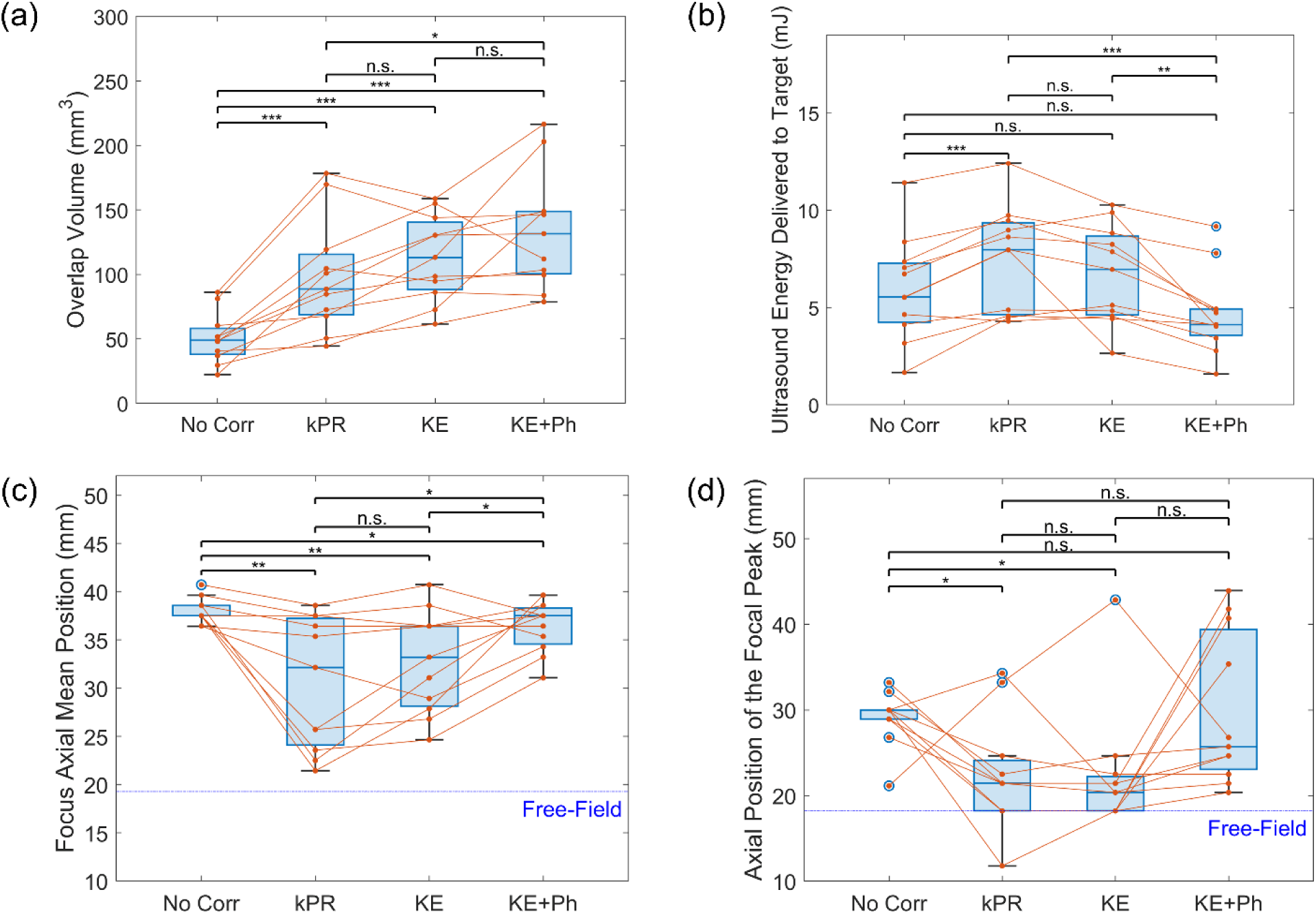
Validation of the efficacy of the proposed phase aberration correction method using the ray-based Kranion software as a reference (*N* = 22. (a) The phase array element activation criterion generated by Kranion, combined with its estimated phase delay values (KE+Ph), achieves the highest overlap volume (*V*_*overlap*_). However, considering overlap volume alone is insufficient for evaluating the focusing performance and ultrasound energy delivery efficiency (as KE+Ph has the largest focus volume); therefore, additional evaluation based on the other three metrics are required. (b) The proposed k-Wave based phase correction method (kPR) effectively increases the ultrasound energy delivered to the target (*E*_*target*_). Compared to KE (using element activation criterion generated by Kranion alone, phase delay estimated by the proposed method), this indicates that the proposed method can accurately estimate the required phase delays for aberration correction and that activating only a subset of array elements (possibly those with smaller incident angles) does not significantly affect energy projection efficiency. (c) The proposed method (kPR) best restores the axial mean position of the focal region (*X*_*focus*_), though KE and KE+Ph also show a certain degree of decrease. (d) Both kPR and KE successfully restore the axial position of the focal peak (*X*_*peak*_), with no statistically significant difference between the two. However, KE+Ph does not adequately improve this metric again. Given such results, the phase delay and element activation criterion estimated by Kranion alone does not sufficiently restore the spatial position of the focus or improve the energy delivery. *Note: Statistical significance was assessed using two-tailed paired t-tests; Key: *p<0.05, **p<0.01, ***p<0.001, n.s.: p≥0.05*.

**Table III.** Quantitative Comparison of Ultrasound Focusing and Energy Delivery Metrics Across Aberration Correction Methods (*N* = 11; No Corr: no correction, kPR: phase delay estimation by the proposed k-Wave based phase-reversal method with all 128 elements activated, KE: phase delay estimation by the proposed method with element activation criterion by Kranion, KE+Ph: both phase delay estimation and element activation criterion by Kranion)

| | $X_{focus}/\text{mm}$ | $Y_{focus}/\text{mm}$ | $Z_{focus}/\text{mm}$ | $d_x/\text{mm}$ | $d_y/\text{mm}$ | $d_z/\text{mm}$ | $X_{peak}/\text{mm}$ | $V_{overlap}/\text{mm}^3$ | Overlap Rate/% | $E_{target}/\text{mJ}$ |
| --- | --- | --- | --- | --- | --- | --- | --- | --- | --- | --- |
| No Corr | 37.99±1.30 | 0.00±0.00 | 0.00±0.00 | 31.36±0.84 | 3.51±0.70 | 3.50±0.50 | 29.10±3.13 | 50.61±19.61 | 2.84±1.13 | 5.96±2.67 |
| kPR | **30.58±6.82** | **0.19±0.43** | **-0.19±0.43** | **29.51±14.11** | **3.51±1.19** | **3.80±1.47** | **22.30±6.57** | 98.27±43.69 | 5.36±2.22 | **7.58±2.68** |
| KE | 32.82±5.30 | 0.29±0.50 | -0.49±0.56 | 36.23±10.92 | 3.60±1.20 | 3.80±1.49 | 22.40±7.10 | 113.16±33.11 | 6.25±2.06 | 6.70±2.52 |
| KE+Ph | 36.33±2.60 | -0.49±1.00 | -0.49±1.00 | 40.62±4.51 | 5.36±1.98 | 5.55±2.19 | 29.81±8.86 | **133.84±45.12** | **7.32±2.33** | 4.69±2.14 |

To further understand if kPR and KE achieved such higher *V*_*overlap*_ through compromising their axial positioning accuracy or focal shape like KE+Ph, we have also compared their *X*_*focus*_ (Figure 6c), *X*_*peak*_ (Figure 6d), *d*_*x*_ (Table III), and *V*_*focus*_. As a result, kPR (*X*_*focus*,*kPR*_ = 30.58±6.82 mm, *p* = 3.53×10^-3^; *X*_*peak*,*kPR*_ = 22.30±6.57 mm, *p* = 2.28×10^-2^) and KE (*X*_*focus*,*kE*_ = 32.82±5.30 mm, *p* = 4.89×10^-3^; *X*_*peak*,*kE*_ = 22.40±7.10 mm, *p* = 4.39×10^-2^) have significantly smaller values of *X*_*focus*_ and *X*_*peak*_ compared to that of No Corr (*X*_*focus*,*NoCorr*_ = 37.99±1.30 mm; *X*_*peak*,*NoCorr*_ = 29.10±3.13 mm). Meanwhile, although kPR (29.51±14.11 mm, *p* = 0.67) has a smaller *d*_*x*_ compared to No Corr (31.36±0.84 mm), and KE (36.23±10.92 mm, *p* = 0.17) has a larger *d*_*x*_ compared to No Corr, there is no significant difference for this metric between kPR and No Corr as well as between KE and No Corr. Furthermore, kPR (285.24±201.89 mm^3^, *p* = 0.61) has a smaller *V*_*focus*_ compared to No Corr (316.44±16.69 mm^3^) while KE (359.82±221.54 mm^3^, *p* = 0.52) has a larger *V*_*focus*_ compared to No Corr, but similarly, there is no significant difference for this metric. Therefore, kPR and KE can both effectively improve the axial positioning accuracy and targeting specificity of tFUS while showing no significant compromise in terms of the shape of focus.

Furthermore, *E*_*target*_ (Figure 6b) also shows that using Kranion (i.e., KE and KE+Ph) cannot well address the skull-induced aberration. This potential limitation of Kranion arises from KE+Ph (4.69±2.14 mJ) having a significantly lower *E*_*target*_ compared to KE (6.70±2.52 mJ, *p* = 1.75×10^-3^), while there is no significant difference between KE+Ph and No Corr (5.96±2.67 mJ, *p* = 5.68×10^-2^) or between KE (*p* = 0.21) and No Corr. However, kPR (7.58±2.68 mJ, *p* = 2.14×10^-4^) can still offer a significantly higher *E*_*target*_compared to No Corr, aligning with the results for a larger subject group shown in previous subsections.

Therefore, the proposed phase-reversal-based aberration correction method outperforms the ray-based method implemented in Kranion in terms of axial positioning, targeting specificity, and ultrasound energy delivery, without compromising the focal shape or requiring additional estimation of element activation criterion.

## 4. Discussion

In this study, we have introduced and validated a phase-reversal based aberration correction method specifically designed for transcranial focused ultrasound (tFUS) neuromodulation applications, targeting the human left visual cortex V5 (V5L). Our k-Wave simulation results significantly enhance both the spatial positioning accuracy and the efficiency of ultrasound energy delivery through a 128-element random phased-array transducer, thus addressing major limitations inherent in existing transcranial neuromodulation techniques using 1D-array transducer or large phased-array transducer.

The primary finding of this study, i.e., the substantial improvement in targeting accuracy, addresses critical challenges faced in tFUS applications, specifically the skull-induced aberration effects. The skull’s complex geometry and acoustic properties can distort and shift the ultrasound focus, thus limiting the clinical and research utility of tFUS. By employing individualized subject-specific head models and implementing a phase-reversal method, we achieved a 98.70% increase in the overlap volume (*V*_*overlap*_) between the ultrasound focus and V5L, and a significant reduction in axial positioning errors (14.36% improvement in the axial mean position of the focal region, *X*_*focus*_, and 21.53% improvement in focal peak axial position, *X*_*peak*_). This is crucial because precise targeting of brain regions such as V5L can further enhance the specificity and efficacy of neuromodulation protocols, particularly for therapeutic interventions and tFUS-brain-computer interface (tFUS-BCI) studies. Furthermore, improved targeting helps ultrasound energy delivery to the target region. From our results, the total ultrasound energy delivered to V5L (*E*_*target*_) significantly increased by 17.58%, which proves that the proposed aberration correction method can effectively project the necessary energy for neuromodulatory effects thermally or mechanically to the target region.

Notably, we found no significant difference in the axial diameter of the focal region (*d*_*x*_) on a group level as shown in Table I (*N* = 22; 31.38±3.22 mm before correction and 32.19±12.68 mm after correction, *p* = 0.77). This implies that the proposed aberration correction method may not be helpful for improving the axial spatial resolution of tFUS for certain cases, though we have observed a decrease of *d*_*x*_ on several subjects (*N* = 8) while there is no change on another few subjects (*N* = 2). However, when considering that tFUS (even prior to correction) has already demonstrated exceptional spatial confinement compared to other non-invasive neuromodulation methods like tDCS and TMS [31], tFUS is still more suitable for targeted neural interventions even with no further improvement on its axial spatial resolution. The aberration correction method proposed in this study further enhances this inherent advantage (i.e., high spatial resolution) of tFUS by improving its axial positioning accuracy and energy delivery efficiency, enabling even more precise and effective modulation of neural circuits.

When comparing to our additional simulation results with phase delay or element activation criterion estimated by Kranion, it may still be difficult to explicitly understand why using Kranion can generate the largest *V*_*overlap*_ but only achieves the worst in terms of energy delivery (*E*_*target*_). To further clarify this, we investigated what caused the significant difference of *E*_*target*_ between KE and KE+Ph as there is no group-level difference between kPR and KE (Figure 6). First, we need to combine Equation 9 and Equation 10 to see how *E*_*target*_ was calculated:

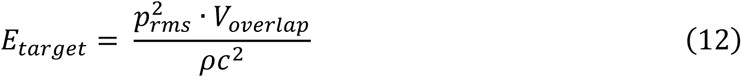

In Equation 12, *ρ* and *c* are constants defined by the medium density and speed of sound in the medium, respectively. So, only *p*_*rms*_ and *V*_*overlap*_ determine how much energy can be delivered to the target region. Through additional calculations, KE (383.75±62.81 kPa) has a significantly larger *p*_*rms*_ compared to KE+Ph (296.37±68.17 kPa, *p* = 9.91×10^-4^), while KE has a similar *V*_*overlap*_ compared to KE+Ph as mentioned above (*p* = 0.11). This explains why KE+Ph generates a significantly lower *E*_*target*_ even it has the largest *V*_*overlap*_, making it the less effective case in terms of ultrasound energy delivery.

Additionally, further analysis on KE reveals that selectively activating a subgroup of elements can also correct the aberration to a certain degree, i.e., KE shows significant improvements on *V*_*overlap*_ (Figure 6a), *X*_*focus*_ (Figure 6c), and *X*_*peak*_ (Figure 6d). This implies that the other subgroup of elements may have a large enough incident angle that effectively prevents the ultrasound waves generated from such elements to transmit through the skull, i.e., most of the ultrasound energy can be reflected back to the extracranial space. Such findings can also be helpful in future studies on random array transducer design (a smaller array with fewer elements may efficiently produce the same ultrasound focus) as well as in designing more effective algorithms to estimate element activation criterions. More importantly, only the proposed method (kPR) can significantly improve all metrics, aligning with the results above on a larger subject group, implying minimum rewards by estimating the element activation criterion using Kranion (i.e., there is no significant difference between kPR and KE in terms of all four metrics). Such comparisons also underscore the necessity of accurate phase delay estimation provided by the proposed phase-reversal approach using k-Wave, which better accounts for complex acoustic phenomena such as diffraction and scattering.

Despite such strengths of the proposed method, several limitations should also be acknowledged. First, this study primarily employed simulated data based on MR-derived pseudo-CT (pCT) skull models. While this approach minimizes radiation exposure and is practical for patient-specific planning, the accuracy of pCT and the binary models in capturing precise skull density variations may impact real-world generalizability. Meanwhile, the use of a fixed ultrasound frequency (700 kHz) and continuous wave (CW) source signals in this work restricts the direct extrapolation of our findings to other frequencies or source configurations that might be employed experimentally. Moreover, although the k-Wave toolbox is a widely used computational tool for simulating acoustic wave propagation, particularly in complex and heterogeneous media like the skull and brain tissues, when compared to actual measurements, limitations may arise due to inherent assumptions and practical constraints within the simulation environment. In addition, solving the first-order k-space model in a 3D space can be relatively time-consuming just like other numerical methods while GPU acceleration may greatly alleviate this problem. Specifically, we compared the time consumption on a same 128×162×162 3D grid using the same simulation settings discussed in Section *II-C Acoustic Simulation Setup* with different solvers. The kspaceFirstOrder3D solver working on CPU (MATLAB) takes about 58.84 seconds per simulation while the kspaceFirstOrder3DG solver working on single GPU (C++/CUDA) takes only 11.77 seconds, which saves 80% of time per simulation. For our 128-element phase delay estimations, the GPU (C++/CUDA) solver can save around 1 hour on average even compared to the GPU (MATLAB) solver (from around 5.5 hours reduced to around 4.5 hours). Such time consumption can be further reduced by using multi-GPUs [76], or by using a pre-calculated source matrix in our workload (this can further reduce the total computation time to around 3 hours when using the GPU (C++/CUDA) solver).

Therefore, future research should address such limitations by integrating empirical validation through experimental *in-vivo* or *ex-vivo* measurements, possibly combining simulations with empirical transcranial ultrasound field measurements. Additionally, investigating the effectiveness of the proposed approach at varying ultrasound frequencies and transducer configurations would also significantly facilitate clinical translation. Further, extending it to deeper brain targets and different cortical regions (when using the same or different transducers) will expand the applicability and robustness of tFUS neuromodulation or therapeutic applications.

In short, this work represents an advancement in the precision of tFUS neuromodulation, significantly improving focal accuracy and energy delivery efficiency through phase aberration correction. This not only positions tFUS as an even more compelling alternative to other invasive and noninvasive neuromodulation techniques, but also opens the door for more targeted and efficient interventions in neuroscience research and clinical management of neurological disorders. By refining the targeting specificity of transcranial ultrasound, we facilitate the future development of safer, more effective, and highly personalized neuromodulation therapies.

## 5. Conclusion

In this work, we have developed a k-Wave-based phase-reversal aberration correction method tailored for a 128-element random phased-array transducer targeting the human left V5 (V5L) region. This method reliably corrected skull-induced phase aberrations, yielding a 98.70% increase in focus–target overlap volume, a 14.36% reduction in mean axial positioning error, a 21.53% reduction in focal peak axial positioning error, and a 17.6% improvement in delivered energy versus uncorrected simulations, which consistently outperforms the ray-tracing approach in Kranion. These improvements in spatial precision and energy deposition underscore the importance of accurate phase-delay estimation for personalized tFUS neuromodulation and pave the way for reliable non-invasive brain stimulation protocols in humans minimizing off-target effects. Future work is warranted to validate this method experimentally, examine other transducer designs and frequencies, and optimize computational efficiency to enable real-time applications.

## 6. Data Availability Statement

The data that support the findings of this study are provided in the paper. k-Wave simulation codes are available at the following GitHub repository: https://github.com/bfinl/tFUS-Aberration-Correction-kWave

## 7. Conflict of Interest Statement

B.H., K.Y., and J.K. are co-inventors of pending patent applications. Z.L. has no conflict of interest to declare.

## 8. Funding Statement

This work was supported in part by National Institutes of Health grants RF1NS131069, R01NS124564, U18EB029354, T32EB029365, R01NS096761, and R01NS127849-01A1. J.K. was supported in part by the National Science Foundation Graduate Research Fellowship Program grant DGE2140739.

## 9. Ethical Statement

Our study complied with all relevant ethical regulations regarding human research and were reviewed and approved by the Advarra Institutional Review Board (protocol number: STUDY2017_00000426). Subjects were safety screened for MRI eligibility and informed of the potential risks of the study. Subjects who wished to continue gave voluntary informed consent in accordance with the World Medical Association’s Declaration of Helsinki. Subjects were compensated for their time at a rate of $20 per hour. We did not obtain consent to publish subject-identifying information.

## Notes

### Competing Interest Statement

The authors have declared no competing interest.

### Summary of Updates

Updated manuscript format, conclusion section, and data availability statement.

## References

[1] R. Abrams, Electroconvulsive therapy, Oxford University Press, 2002.

[2] K. E. Hoy and P. B. Fitzgerald, “Brain stimulation in psychiatry and its effects on cognition,” Nature Reviews Neurology, vol. 6, no. 5, pp. 267–275, 2010.

[3] W. J. Fry, F. Fry, J. Barnard, R. Krumins, and J. Brennan, “Ultrasonic lesions in the mammalian central nervous system,” Science, vol. 122, no. 3168, pp. 517–518, 1955.

[4] D. Coluccia, J. Fandino, L. Schwyzer, R. O’Gorman, L. Remonda, J. Anon, E. Martin, and B. Werner, “First noninvasive thermal ablation of a brain tumor with mr-guided focused ultrasound,” Journal of therapeutic ultrasound, vol. 2, pp. 1–7, 2014.

[5] W. J. Tyler, Y. Tufail, M. Finsterwald, M. L. Tauchmann, E. J. Olson, and C. Majestic, “Remote excitation of neuronal circuits using low-intensity, low-frequency ultrasound,” PloS one, vol. 3, no. 10, pp. e3511, 2008.

[6] E. Mehic, J. M. Xu, C. J. Caler, N. K. Coulson, C. T. Moritz, and P. D. Mourad, “Increased anatomical specificity of neuromodulation via modulated focused ultrasound,” PloS one, vol. 9, no. 2, pp. e86939, 2014.

[7] C. Liu, K. Yu, X. Niu, and B. He, “Transcranial focused ultrasound enhances sensory discrimination capability through somatosensory cortical excitation,” Ultrasound in medicine & biology, vol. 47, no. 5, pp. 1356–1366, 2021.

[8] K. Yu, C. Liu, X. Niu, and B. He, “Transcranial focused ultrasound neuromodulation of voluntary movement-related cortical activity in humans,” IEEE Transactions on Biomedical Engineering, vol. 68, no. 6, pp. 1923–1931, 2020.

[9] H. Kim, M. Y. Park, S. D. Lee, W. Lee, A. Chiu, and S.-S. Yoo, “Suppression of eeg visual-evoked potentials in rats through neuromodulatory focused ultrasound,” Neuroreport, vol. 26, no. 4, pp. 211–215, 2015.

[10] T. Zhang, N. Pan, Y. Wang, C. Liu, and S. Hu, “Transcranial focused ultrasound neuromodulation: a review of the excitatory and inhibitory effects on brain activity in human and animals,” Frontiers in human neuroscience, vol. 15, pp. 749162, 2021.

[11] K. Yu, X. Niu, E. Krook-Magnuson, and B. He, “Intrinsic functional neuron-type selectivity of transcranial focused ultrasound neuromodulation,” Nature communications, vol. 12, no. 1, p. 2519, 2021.

[12] X. Niu, K. Yu, and B. He, “Transcranial focused ultrasound induces sustained synaptic plasticity in rat hippocampus,” Brain stimulation, vol. 15, no. 2, pp. 352–359, 2022.

[13] S. Ramachandran, X. Niu, K. Yu, and B. He, “Transcranial ultrasound neuromodulation induces neuronal correlation change in the rat somatosensory cortex,” Journal of neural engineering, vol. 19, no. 5, p. 056002, 2022.

[14] L. di Biase, E. Falato, M. L. Caminiti, P. M. Pecoraro, F. Narducci, and V. Di Lazzaro, “Focused ultrasound (fus) for chronic pain management: approved and potential applications,” Neurology research international, vol. 2021, no. 1, pp. 8438498, 2021.

[15] B. W. Badran and X. Peng, “Transcranial focused ultrasound (tfus): A promising noninvasive deep brain stimulation approach for pain,” Neuropsychopharmacology, vol. 49, no. 1, pp. 351–352, 2024.

[16] M. G. Kim, K. Yu, C.-Y. Yeh, R. Fouda, D. Argueta, S. Kiven, Y. Ni, X. Niu, Q. Chen, K. Kim et al., “Low-intensity transcranial focused ultrasound suppresses pain by modulating pain-processing brain circuits,” Blood, vol. 144, no. 10, pp. 1101–1115, 2024.

[17] Y. Meng, K. Hynynen, and N. Lipsman, “Applications of focused ultrasound in the brain: from thermoablation to drug delivery,” Nature Reviews Neurology, vol. 17, no. 1, pp. 7–22, 2021.

[18] X. Liu, S. S. M. Naomi, W. L. Sharon, and E. J. Russell, “The applications of focused ultrasound (FUS) in alzheimer’s disease treatment: a systematic review on both animal and human studies,” Aging and disease, vol. 12, no. 8, pp. 1977, 2021.

[19] J. Vidal-Jove, X. Serres, E. Vlaisavljevich, J. Cannata, A. Duryea, R. Miller, X. Merino, M. Velat, Y. Kam, R. Bolduan et al., “First-in-man histotripsy of hepatic tumors: the theresa trial, a feasibility study,” International Journal of Hyperthermia, vol. 39, no. 1, pp. 1115–1123, 2022.

[20] N. Khanna, D. Gandhi, A. Steven, V. Frenkel, and E. Melhem, “Intracranial applications of mr imaging–guided focused ultrasound,” American Journal of Neuroradiology, vol. 38, no. 3, pp. 426–431, 2017.

[21] M. G. Kim, C.-Y. Yeh, K. Yu, Z. Li, K. Gupta, and B. He, “Analgesic effect of simultaneously targeting multiple pain processing brain circuits in an aged humanized mouse model of chronic pain by transcranial focused ultrasound,” APL bioengineering, vol. 9, no. 1, 2025.

[22] K. Yu, X. Niu, and B. He, “Neuromodulation management of chronic neuropathic pain in the central nervous system,” Advanced Functional Materials, vol. 30, no. 37, p. 1908999, 2020.

[23] J. Hubble, K. Busenbark, S. Wilkinson, R. Penn, K. Lyons, and W. Koller, “Deep brain stimulation for essential tremor,” Neurology, vol. 46, no. 4, pp. 1150–1153, 1996.

[24] G. Deuschl, C. Schade-Brittinger, P. Krack, J. Volkmann, H. Schafer, K. Botzel, C. Daniels, A. Deutschlander, U. Dillmann, W. Eisner et al., “A randomized trial of deep-brain stimulation for parkinson’s disease,” New England Journal of Medicine, vol. 355, no. 9, pp. 896–908, 2006.

[25] M. Hodaie, R. A. Wennberg, J. O. Dostrovsky, and A. M. Lozano, “Chronic anterior thalamus stimulation for intractable epilepsy,” Epilepsia, vol. 43, no. 6, pp. 603–608, 2002.

[26] K. Ishibashi, K. Shimada, T. Kawato, S. Kaji, M. Maeno, S. Sato, and K. Ito, “Inhibitory effects of low-energy pulsed ultrasonic stimulation on cell surface protein antigen c through heat shock proteins groel and dnak in streptococcus mutans,” Applied and Environmental Microbiology, vol. 76, no. 3, pp. 751–756, 2010.

[27] A.-H. Javadi, A. Beyko, V. Walsh, and R. Kanai, “Transcranial direct current stimulation of the motor cortex biases action choice in a perceptual decision task,” Journal of cognitive neuroscience, vol. 27, no. 11, pp. 2174–2185, 2015.

[28] L.M. Koponen and A.V. Peterchev, “Transcranial Magnetic Stimulation: Principles and Applications,” In: B. He (ed), Neural engineering. Springer, pp. 245–270, 2020.

[29] Z.-D. Deng, S. H. Lisanby, and A. V. Peterchev, “Electric field depth–focality tradeoff in transcranial magnetic stimulation: simulation comparison of 50 coil designs,” Brain stimulation, vol. 6, no. 1, pp. 1–13, 2013.

[30] ED. Edwards, M. Cortes, A. Datta, P. Minhas, E. M. Wassermann, and M. Bikson, “Physiological and modeling evidence for focal transcranial electrical brain stimulation in humans: a basis for high-definition tdcs,” Neuroimage, vol. 74, pp. 266–275, 2013.

[31] A. Bystritsky, A. S. Korb, P. K. Douglas, M. S. Cohen, W. P. Melega, A. P. Mulgaonkar, A. DeSalles, B.-K. Min, and S.-S. Yoo, “A review of low-intensity focused ultrasound pulsation,” Brain stimulation, vol. 4, no. 3, pp. 125–136, 2011.

[32] D. White, J. Clark, J. Chesebrough, M. White, and J. Campbell, “Effect of the skull in degrading the display of echoencephalographic b and c scans,” The Journal of the Acoustical Society of America, vol. 44, no. 5, pp. 1339–1345, 1968.

[33] J.-F. Aubry and M. Tanter, “Mr-guided transcranial focused ultrasound,” Therapeutic ultrasound, pp. 97–111, 2016.

[34] M. Wang, Z. Xu, and B. Cheng, “Systematic review of phase aberration correction algorithms for transcranial focused ultrasound,” Iradiology, vol. 3, no. 1, pp. 26–46, 2025.

[35] G. T. Clement and K. Hynynen, “Micro-receiver guided transcranial beam steering,” IEEE transactions on ultrasonics, ferroelectrics, and frequency control, vol. 49, no. 4, pp. 447–453, 2002.

[36] T. Riis, D. Feldman, B. Mickey, and J. Kubanek, “Controlled noninvasive modulation of deep brain regions in humans,” Communications Engineering, vol. 3, no. 1, pp. 13, 2024.

[37] T. Riis, M. Wilson, and J. Kubanek, “Controlled delivery of ultrasound through the head for effective and safe therapies of the brain,” bioRxiv, pp. 2022–12, 2022.

[38] F. Sammartino, D. W. Beam, J. Snell, and V. Krishna, “Kranion, an open-source environment for planning transcranial focused ultrasound surgery,” Journal of neurosurgery, vol. 132, no. 4, pp. 1249–1255, 2019.

[39] N. Lu, T. L. Hall, J. R. Sukovich, S. W. Choi, J. Snell, N. McDannold, and Z. Xu, “Two-step aberration correction: application to transcranial histotripsy,” Physics in Medicine & Biology, vol. 67, no. 12, pp. 125009, 2022.

[40] C. Jin, D. Moore, J. Snell, and D.-G. Paeng, “An open-source phase correction toolkit for transcranial focused ultrasound,” BMC biomedical engineering, vol. 2, pp. 1–11, 2020.

[41] B. Li, L. Xie, C. Liu, K. Xu, Y. Zhan, D. Ta et al., “Ray theory-based compounded plane wave ultrasound imaging for aberration corrected transcranial imaging: Phantom experiments and simulations,” Ultrasonics, vol. 135, pp. 107124, 2023.

[42] J. L. Robertson, B. T. Cox, J. Jaros, and B. E. Treeby, “Accurate simulation of transcranial ultrasound propagation for ultrasonic neuromodulation and stimulation,” The Journal of the Acoustical Society of America, vol. 141, no. 3, pp. 1726–1738, 2017.

[43] Y. Jing, F. C. Meral, and G. T. Clement, “Time-reversal transcranial ultrasound beam focusing using a k-space method,” Physics in Medicine & Biology, vol. 57, no. 4, pp. 901, 2012.

[44] J. Gu and Y. Jing, “Modeling of wave propagation for medical ultrasound: a review,” IEEE transactions on ultrasonics, ferroelectrics, and frequency control, vol. 62, no. 11, pp. 1979–1992, 2015.

[45] K. Yee, “Numerical solution of initial boundary value problems involving maxwell’s equations in isotropic media,” IEEE Transactions on antennas and propagation, vol. 14, no. 3, pp. 302–307, 1966.

[46] T. D. Mast, L. P. Souriau, D.-L. Liu, M. Tabei, A. I. Nachman, and R. C. Waag, “A k-space method for large-scale models of wave propagation in tissue,” IEEE transactions on ultrasonics, ferroelectrics, and frequency control, vol. 48, no. 2, pp. 341–354, 2001.

[47] B. E. Treeby and B. T. Cox, “k-wave: Matlab toolbox for the simulation and reconstruction of photoacoustic wave fields,” Journal of biomedical optics, vol. 15, no. 2, pp. 021314–021314, 2010.

[48] C. Angla, B. Larrat, J.-L. Gennisson, and S. Chatillon, “Transcranial ultrasound simulations: A review,” Medical Physics, vol. 50, no. 2, pp. 1051–1072, 2023.

[49] S. A. Leung, D. Moore, Y. Gilbo, J. Snell, T. D. Webb, C. H. Meyer, G. W. Miller, P. Ghanouni, and K. Butts Pauly, “Comparison between mr and ct imaging used to correct for skull-induced phase aberrations during transcranial focused ultrasound,” Scientific reports, vol. 12, no. 1, pp. 13407, 2022.

[50] L. Demi, “Practical guide to ultrasound beam forming: Beam pattern and image reconstruction analysis,” Applied Sciences, vol. 8, no. 9, pp. 1544, 2018.

[51] O. T. Von Ramm and S. W. Smith, “Beam steering with linear arrays,” IEEE transactions on biomedical engineering, no. 8, pp. 438–452, 1983.

[52] J. Hand, A. Shaw, N. Sadhoo, S. Rajagopal, R. Dickinson, and L. Gavrilov, “A random phased array device for delivery of high intensity focused ultrasound,” Physics in Medicine & Biology, vol. 54, no. 19, pp. 5675, 2009.

[53] G. T. Clement, J. White, and K. Hynynen, “Investigation of a large-area phased array for focused ultrasound surgery through the skull,” Physics in Medicine & Biology, vol. 45, no. 4, pp. 1071, 2000.

[54] L. Marsac, D. Chauvet, R. La Greca, A.-L. Boch, K. Chaumoitre, M. Tanter, and J.-F. Aubry, “Ex vivo optimisation of a heterogeneous speed of sound model of the human skull for noninvasive transcranial focused ultrasound at 1 mhz,” International Journal of Hyperthermia, vol. 33, no. 6, pp. 635–645, 2017.

[55] A. Javid, S. Ilham, and M. Kiani, “A review of ultrasound neuromodulation technologies,” IEEE transactions on biomedical circuits and systems, vol. 17, no. 5, pp. 1084–1096, 2023.

[56] J. Kosnoff, K. Yu, C. Liu, and B. He, “Transcranial focused ultrasound to v5 enhances human visual motion brain-computer interface by modulating feature-based attention,” Nature Communications, vol. 15, no. 1, pp. 4382, 2024.

[57] J. C. Mazziotta, A. W. Toga, A. Evans, P. Fox, J. Lancaster et al., “A probabilistic atlas of the human brain: theory and rationale for its development,” Neuroimage, vol. 2, no. 2, pp. 89–101, 1995.

[58] J. Mazziotta, A. Toga, A. Evans, P. Fox, J. Lancaster, K. Zilles, R. Woods, T. Paus, G. Simpson, B. Pike et al., “A probabilistic atlas and reference system for the human brain: International consortium for brain mapping (icbm),” Philosophical Transactions of the Royal Society of London. Series B: Biological Sciences, vol. 356, no. 1412, pp. 1293– 1322, 2001.

[59] J. Mazziotta, A. Toga, A. Evans, P. Fox, J. Lancaster, K. Zilles, R. Woods, T. Paus, G. Simpson, B. Pike et al., “A four-dimensional probabilistic atlas of the human brain,” Journal of the American Medical Informatics Association, vol. 8, no. 5, pp. 401–430, 2001.

[60] R. T. Born and D. C. Bradley, “Structure and function of visual area mt,” Annu. Rev. Neurosci., vol. 28, no. 1, pp. 157–189, 2005.

[61] M. Jenkinson, C. F. Beckmann, T. E. Behrens, M. W. Woolrich, and S. M. Smith, “Fsl,” Neuroimage, vol. 62, no. 2, pp. 782–790, 2012.

[62] S. M. Smith, “Fast robust automated brain extraction,” Human brain mapping, vol. 17, no. 3, pp. 143–155, 2002.

[63] M. Jenkinson, P. Bannister, M. Brady, and S. Smith, “Improved optimization for the robust and accurate linear registration and motion correction of brain images,” Neuroimage, vol. 17, no. 2, pp. 825–841, 2002.

[64] J. L. Andersson, M. Jenkinson, S. Smith et al., “Nonlinear registration, aka spatial normalisation fmrib technical report tr07ja2,” FMRIB Analysis Group of the University of Oxford, vol. 2, no. 1, 2007.

[65] S. B. Eickhoff, K. E. Stephan, H. Mohlberg, C. Grefkes, G. R. Fink, K. Amunts, and K. Zilles, “A new spm toolbox for combining probabilistic cytoarchitectonic maps and functional imaging data,” Neuroimage, vol. 25, no. 4, pp. 1325–1335, 2005.

[66] S. N. Yaakub, T. A. White, E. Kerfoot, L. Verhagen, A. Hammers, and E. F. Fouragnan, “Pseudo-cts from t1-weighted mri for planning of low-intensity transcranial focused ultrasound neuromodulation: An open-source tool,” Brain Stimulation: Basic, Translational, and Clinical Research in Neuromodulation, vol. 16, no. 1, pp. 75–78, 2023.

[67] J. A. Cain, S. Visagan, and M. M. Monti, “Smartfus: surrogate model of attenuation and refraction in transcranial focused ultrasound,” Plos one, vol. 17, no. 10, pp. e0264101, 2022.

[68] J. K. Mueller, L. Ai, P. Bansal, and W. Legon, “Numerical evaluation of the skull for human neuromodulation with transcranial focused ultrasound,” Journal of neural engineering, vol. 14, no. 6, pp. 066012, 2017.

[69] J. Blackmore, S. Shrivastava, J. Sallet, C. R. Butler, and R. O. Cleveland, “Ultrasound neuromodulation: a review of results, mechanisms and safety,” Ultrasound in medicine & biology, vol. 45, no. 7, pp. 1509–1536, 2019.

[70] Y. Huang, P. Wen, B. Song, and Y. Li, “Numerical investigation of the energy distribution of low-intensity transcranial focused ultrasound neuromodulation for hippocampus,” Ultrasonics, vol. 124, pp. 106724, 2022.

[71] J. O. Smith III, Physical audio signal processing: For virtual musical instruments and audio effects, W3K Publishing, 2010, [Online]. Available: https://ccrma.stanford.edu/~jos/pasp/AcousticEnergyDensity.html

[72] Wikipedia contributors, “Sound energy — Wikipedia, the free encyclopedia,” 2025, [Online]. Available: https://en.wikipedia.org/w/index.php?title=Sound\energy\&oldid=1271867582

[73] A. D. Pierce, Acoustics: an introduction to its physical principles and applications, Springer, 2019.

[74] D. Raboud and L. Westover, Sound and Acoustics: Sound Pressure Measurements, Engineeringat Alberta — Open Educational Resources, [Online]. Available: https://engcourses-uofa.ca/books/vibrations-and-sound/12-sound-and-acoustics/sound-pressure-measurements/

[75] MATLAB version 24.2.0.2863752 (R2024b) Update 5, The Mathworks, Inc., Natick, Massachusetts, 2024.

[76] B. Treeby, F. Vaverka, and J. Jaros, “Performance and accuracy analysis of nonlinear k-wave simulations using local domain decomposition with an 8-gpu server,” in Proceedings of Meetings on Acoustics. AIP Publishing, 2018, vol. 34.

